# Two-photon frequency division multiplexing for functional *in vivo* imaging: a feasibility study

**DOI:** 10.1101/405118

**Authors:** Dmitri Tsyboulski, Natalia Orlova, Peter Ledochowitsch, Peter Saggau

**Affiliations:** Allen Institute, 615 Westlake Ave N, Seattle, WA 98109

## Abstract

Recently, we presented a new approach to create high-speed amplitude modulation of femtosecond laser pulses and tag multiple excitation beams with specific modulation frequencies. In this work, we discuss the utility of this method to record calcium signals in brain tissue with two-photon frequency-division multiplexing (2P-FDM) microscopy. While frequency-multiplexed imaging appears slightly inferior in terms of image quality as compared to conventional two-photon laser scanning microscopy due to shot noise-induced cross-talk between frequency channels, applying this technique to record average signals from regions of interest (ROI) such as neuronal cell bodies was found to be promising. We use phase information associated with each pixel or waveform within a selected ROI to phase-align and recombine the signals into one extended amplitude-modulated waveform. This procedure narrows the frequency detection window, effectively decreasing noise contributions from other frequency channels. Using theoretical analysis, numerical simulations, and *in vitro* imaging we demonstrate a reduction of cross-talk by more than an order of magnitude and predict the usefulness of 2P-FDM for functional studies of brain activity.

## 1. Introduction

Two-photon laser scanning microscopy (TPLSM) [1] has become a standard tool for functional imaging of neural activity with sub-cellular resolution. Conventional TPLSM can achieve data acquisition rates up to 50 frames per second [2, 3], sufficient to monitor calcium–associated fluorescence transients within one or several imaging planes with satisfactory temporal resolution. Although highly desired, simultaneous functional recordings from multiple sites in large ensembles of neurons labeled with calcium- or voltage-sensitive indicators remain challenging due to limitations in TPLSM scan speed and data acquisition rate [2, 4]. A variety of experimental techniques have emerged, designed to improve the speed of imaging a larger area or a volume. These techniques can be generally assigned to several broad categories: those increasing the sweep rate at which a focused beam moves through the volume [5], those involving engineered excitation point spread functions to increase the individual voxel size in a volume [6–8], and those restricting the imaging volume to selected regions of interest (ROI) [9–11]. For example, several reports demonstrated functional imaging of cell activity within (0.5 mm)^3^ at the rate of several Hz [6, 7]. Nevertheless, emergence of new imaging systems with significantly larger field-of-view (FOV) and more than an order of magnitude increase in the accessible imaging volume [3, 12] again raise the issue of limited data acquisition rates in TPLSM.

To address emerging needs of large FOV systems, a separate class of techniques may be employed. These methods are focused on multiplexed image acquisition, where the sample is scanned with multiple excitation beams and the total emission is recorded with a single detector [12–16]. One approach, termed “spatiotemporal multiplexing,” utilized multiple excitation pulse trains delayed in time relative to each other and separated the corresponding emission signals based on their arrival time at a detector [12, 13, 16]. The useful number of acquisition channels was ultimately limited by the laser pulse repetition rate and the fluorescence lifetime of the fluorophore, leaving room for only a few multiplexed channels with commonly used commercial 80 MHz femtosecond pulsed lasers.

Recently we introduced a new two-photon frequency-division multiplexing (2P-FDM) microscopy method to create amplitude modulation of femtosecond laser pulses in the MHz range, based on interference of periodically phase-shifted femtosecond pulse trains, and demonstrated frequency-multiplexed two-photon imaging in test samples [17]. The major limiting factor of this approach is shot noise, with its average spectral density uniformly distributed across the detection frequency bandwidth, therefore, inducing cross-talk between individual frequency channels. While it has been predicted that the signal-to-noise ratios (SNR) in 2P-FDM and conventional TPLSM images are similar, given the same data acquisition rate and the same average emission intensity per channel [17], 2P-FDM data are also contaminated with cross-talk related ghost images from other active channels. At high scanning speeds provided by resonant galvanometer (RG) scanners, individual pixel dwell times become less than 100 ns. Together with low emission intensities, this may result in unacceptably low contrast between individual frequency channels in 2P-FDM. In this work, we analyze the performance of 2P-FDM in more detail and demonstrate its utility for recording calcium signals *in vivo*. Most importantly, we propose a novel method of incorporating phase information to the analysis of functional signals that can reduce crosstalk between frequency-encoded imaging channels by more than an order of magnitude.

## 2. Methods

High-speed amplitude modulation of multiple excitation beams derived from a single mode-locked Ti:Sapphire laser (Chameleon Ultra II, Coherent) was created by interfering femtosecond laser pulses at the wavelength of 920 nm with a periodic phase shift produced by acousto-optic deflectors (AODs, DTSX-A12-800.1000, frequency range 72-113 MHz, AA Opto-electronic) using a custom-built Mach-Zehnder interferometer [17]. The AODs were controlled by frequencies from a 4-channel direct digital synthesizer board (AD9959, Analog Devices). Imaging of tissue slices was performed on a custom-built TPLSM system, equipped with XY galvanometer scanners (6240H, Cambridge Technology), and a water immersion objective lens (25× XLPLAN N, 1.05 NA, Olympus). Emission within a 500-700 nm spectral window was detected by a hybrid avalanche photodetector (R11322U-40, Hamamatsu), connected to a pre-amplifier (DHCPA100, 200 MHz bandwidth, FEMTO), and sampled with a streaming waveform digitizer at 500 MSamples/s (ATS9350, AlazarTech). Galvo-scanners were controlled with two DAQ boards (PCIe-6353, National Instruments), which were programmed to also trigger the waveform acquisition of each scan line. The frame rate was limited to ~ 0.4 Hz by the speed of the galvanometer scanners, which could provide a maximum line scan rate of 200 Hz.

A diagram of the experimental setup is shown in Fig. 1. The femtosecond laser beam is expanded to 9 mm in diameter and directed to the interferometer input. After beam-splitting, the matching pair of beams are directed to the first AOD driven by a single frequency *f* (in this case *f* = 94 MHz) from opposite sides to obtain two diffracted beams with opposite Doppler frequency shifts ±*f*. The diffracted beams are passed through a pair of adjustable delay lines and are diverted to the upper level of the interferometer with periscopic mirrors. Next, the two beams are directed from opposite sides to the second AOD in a similar manner to obtain diffracted beams with ±*f_i_* Doppler frequency shift. The second AOD can be simultaneously driven with multiple frequencies to generate two fans of beams. It is essential to direct the incoming beams such that positive and negative Doppler shifts in the interfering beams are subtracted from each other (see Fig. 1), otherwise, the resulting beam modulation frequency will appear in a range > 300 MHz, far exceeding the current 80 MHz laser pulse rate. After recombining the two beams at the interferometer output, the frequency shift between them becomes |2 ·(*f* – *f_i_*)|, which defines the modulation frequency of the excitation beam. To obtain the modulation frequencies of 1, 5 and 7 MHz used in this work, AOD1 and AOD2 were driven at 94 MHz and a combination of 90.5, 93.5 and 96.5 MHz frequencies, respectively. In general, AODs produce spatial and temporal dispersion of femtosecond laser pulses, and it is essential to correct for these detrimental effects in two-photon imaging. We utilized a previously introduced method to simultaneously compensate for both types of dispersion using a pair of AODs [18]. In Fig. 1, the spatial dispersion of the diffracted beams is illustrated with a color gradient in the diffracted beams after the first AOD. This gradient disappears after the beams pass through the second AOD, indicating that the diverging beams become collimated again. The path length between the AODs defines the magnitude of group delay dispersion compensation, and can be adjusted to maximize the two-photon fluorescence signal. Half-wave plates in the experimental setup control input beam polarization to AODs and ultrafast broadband beam splitters (UBB), which both are polarization-sensitive. Because AODs rotate the output beam polarization by 90°, it is possible to decouple leaving and entering beams with polarizing beam splitters (PBS). After the second AOD both diffracted fans of beams are recombined at the interferometer output to create multiple amplitude-modulated excitation beams. The remaining imaging setup represents a custom-built TPLSM system, where the first 1:1 pupil relay couples AOD2 to the first scanner. The AODs provide limited range of diffraction angles, a total of 2.3° across the 40 MHz frequency range. To fill the back aperture of the objective, the diffracted beams are expanded 1:2 by the scan lens – tube lens relay. In this configuration, AOD2 provides a lateral beam displacement of ~3.6 μm/MHz in the image plane. Note that modulation frequencies are defined by both AODs as described earlier. The intensities of the diffracted beams depend on the amplitude of the corresponding driving frequencies, which allows to adjust the power of individual beams. The detailed performance characterization of the imaging setup is outside of the scope of this manuscript, and will be discussed elsewhere.

**Fig. 1.**
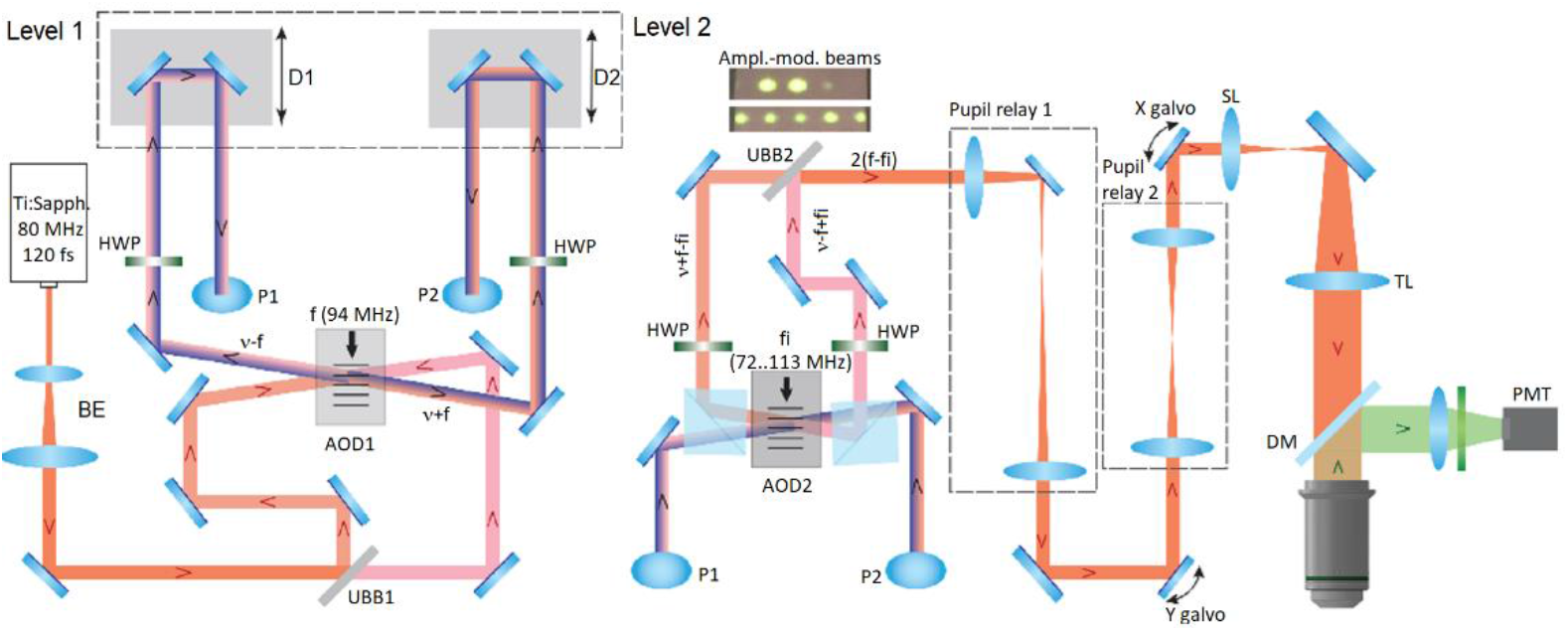
A diagram of the custom Mach-Zehnder interferometric setup to create multiple amplitude-modulated femtosecond laser beams, coupled to the custom-built TPLSM system. BE is beam expander, UBB is 50:50 ultrafast beam splitter, AOD is acousto-optical deflector, HWP is half-wave plate, D is delay line, P is periscopic mirror, PBS is polarizing beam splitter, v is optical frequency, f and f_i_ are acoustic frequencies, SL is scan lens, TL is tube lens, DM is dichroic mirror. The inserted image shows 2 and 5 amplitude-modulated beams at the interferometer output visualized with a NIR detector card.

Custom MATLAB (MathWorks) routines were used for numerical modeling and analysis of amplitude-modulated emission signals. Simulated waveforms were synthesized based on the expected value of a fluorescence signal using Poisson statistics. In simulations, a 160 MHz sampling rate (SR) was selected for convenience, to reduce the amount of data and to obtain sufficient spectral resolution. Note, in real experiments the waveform SR is defined by the laser repetition rate, which also defines the range of usable signal modulation frequencies. Linear combinations of computer-generated waveforms were analyzed in MATLAB using the built-in FFT function to estimate amplitude or power spectrum averages at a given frequency. Error bars in all images correspond to one standard deviation computed from respective datasets.

To demonstrate levels of calcium signals in units of photons/pulse, we calibrated another TPLSM system [3] (Multiphoton Mesoscope, Thorlabs), used in actual *in vivo* imaging experiments. In this system, fluorescence signals were detected using a photomultiplier (HC11706-40, Hamamatsu) coupled to a current amplifier (HCA-400M-5K-C, FEMTO) and a 50 MHz low-pass filter (Mini-Circuits). Waveforms showing single photon detection events were recorded with an oscilloscope (204MXi-A, LeCroy) at 1 GS/s SR without low-pass filtering. Spontaneous neuronal activity was recorded in a Slc17a7-IRES2-Cre mouse, in compliance with the Allen Institute’s animal imaging protocols.

## 3. Cross-talk analysis in frequency-multiplexed two-photon imaging

Measurements of stochastic quantum events such as fluorescence emission are governed by Poisson statistics [19]. At constant excitation intensity, recorded emission signals from a stable and uniform source sampled over discrete time intervals will vary randomly, a phenomenon known as shot noise or Poisson noise, while the ensemble’s variance and mean will remain equal given sufficiently large sample size. For each excitation event *i* at time *t* the detected emission signal *F_i_*(*t*) can be tentatively separated into the sum of an average or expected signal *S_0_* and a purely random (stochastic) noise components *R_i_*(*t*). We may write the following set of equations:

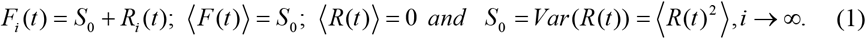

If the expected signal *S*(*t*) varies periodically in time and only its average remains constant, the relationship between the ensemble mean 〈*S*(*t*)〉 and variance of its noise component *R*(*t*) remains the same. Since fluorescence detection events are ergodic and independent, Eq. (1) is valid for the equally probable subsets of samples within time series corresponding to the same expected value. One may recall that the ensemble’s variance is given as the sum of variances of independent components, and the ensemble’s expected value is the weighted sum of expected values of its components. Therefore, one may derive the following expression:

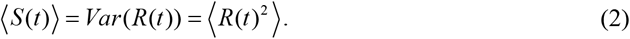

A periodically modulated excitation intensity *I*(*t*) varying from 0 to *I_0_* can be expressed as:

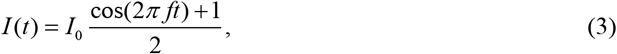

where *I_0_* is the maximum excitation intensity, and *f* is the modulation frequency. The corresponding two-photon emission signal *S*(*t*) is obtained by trigonometric rearrangement from *I^2^*(*t*):

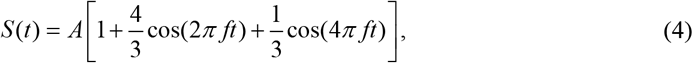

where *A* is the average of the amplitude-modulated emission signal. Here we choose to express *A* in units of photons/sample, which will be useful in the following discussion. Noise variance is related to the power spectrum of a signal. For instance, in the case of discrete measurement of a time series signal, and in the specific case considering our noise component *R*(*t*), Parseval’s theorem states:

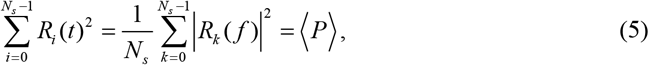

where *R_k_*(*f*) is a discrete Fourier transform (DFT) complex variable corresponding to a frequency *f, N_s_*, is the number of recorded time samples, and 〈*P*〉 is the average of the noise power spectrum. The left part of Eq. (5) equals the sum of the corresponding average values 〈*R_i_*(*t*)^2^〉, and combining with Eq. (2) we obtain:

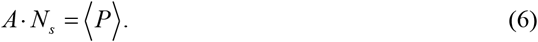

In the case of multiple emission sources, provided they are statistically independent, Eq. (5) can be written as:

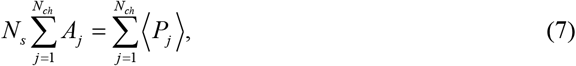

where *N_ch_* represents the number of contributing signals or channels. Eqs. (6) and (7) straightforwardly connect the total average emission signal to the power spectrum average of a corresponding noise time series. To compute an average amplitude spectrum estimate, we recall that DFT values, both real and imaginary, are the weighted sum of random noise variables. With a sufficient number of measurements, these values represent a normal 2D distribution in complex space. One can derive probability distributions of |*R_k_*(*f*)|^2^ and |*R_k_*(*f*)|, which originate from the same normal distribution, and demonstrate that:

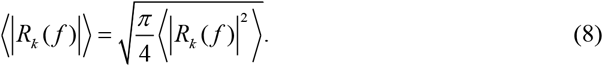

Combining Eqs. (6)–(8), we obtain a background level estimate in the DFT spectrum when multiple noisy signals are added:

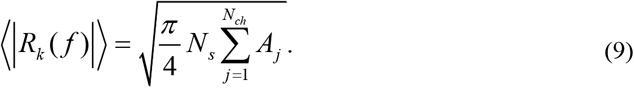

As follows from the definition of DFT (only half of a DFT spectrum is unique), a single harmonic with frequency *f* and amplitude 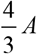, corresponding to the noiseless two-photon signal (see Eq. (2)), has a DFT amplitude of:

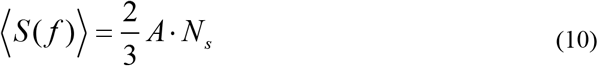

The signal-to-background ratio (SBR) estimate, which is the ratio of the DFT amplitude of a noiseless harmonic and the average background created by noise in all contributing signals, equals:

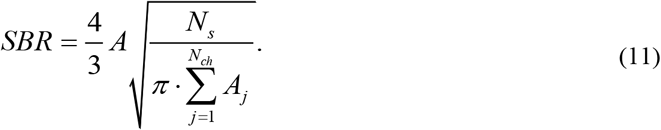

We can see that the SBR increases with the square root of the number of measurements and decreases with the square root of the sum of average emission intensities in each amplitude-modulated channel.

## 4. Numerical simulations comparing 2P-FDM and TPLSM

To demonstrate the validity of the analysis above, we performed numerical simulations of amplitude-modulated signals in the presence of shot noise and estimated signal and cross-talk values for a given number of samples and contributing channels. We synthesized 400 amplitude-modulated waveforms, each containing 8,000 samples for each set of input parameters, which included modulation frequencies *f_i_* = 2*i*-1 MHz, *i*∈[1.10], and average fluorescence signal intensity of 0.5 photons/time sample. Figure 2(a) shows simulated waveforms from a single channel modulated at 7 MHz, and a sum signals from 10 channels. The average amplitude of each waveform is 0.5 photons/sample, corresponding to parameter *A* in Eq. (4). Figure 2(b) shows Fourier spectra of 10 combined waveforms containing 160 and 8,000 samples. The contrast between signals and the background is markedly enhanced with the increased number of samples. Plotting the DFT signal 〈|*S*(7*MHz*)|〉 and background 〈|*R_k_*(21*MHz*)|〉 amplitudes as a function of *N_s_* computed from 400 simulated traces reveals the trends predicted by Eqs. (9) and (10) (Fig. 2(c)). Solid lines show theoretical predictions which are in excellent agreement with our simulation results.

**Fig. 2.**
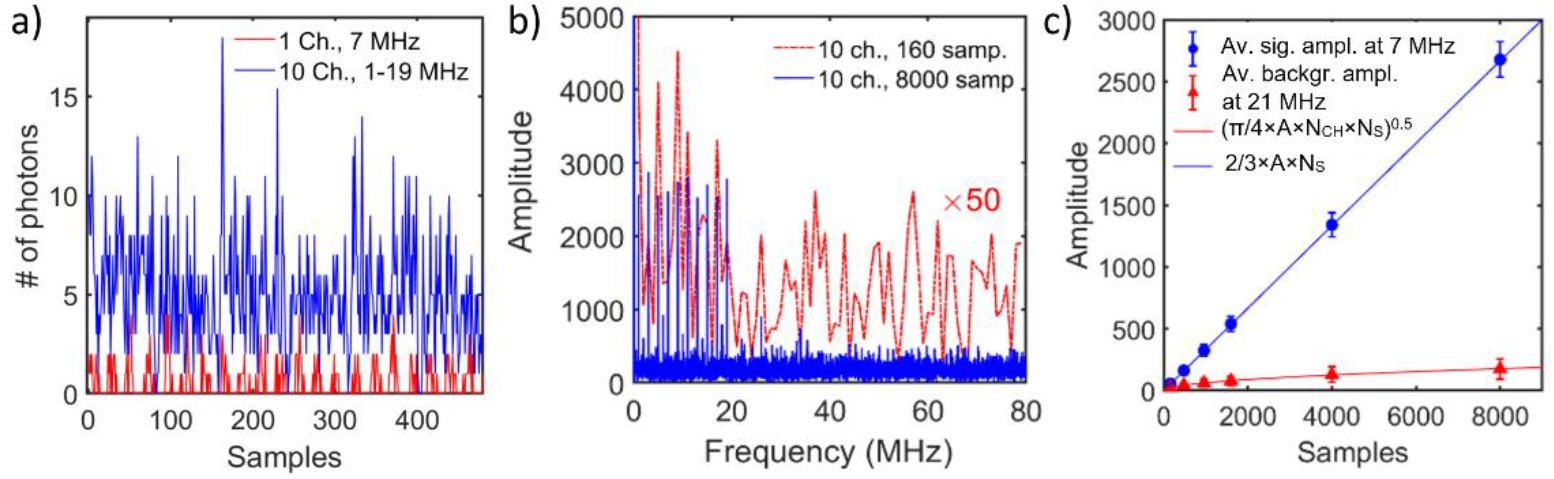
(a) Computer-generated waveforms from 1 and 10 simultaneous channels, amplitude-modulated at 7 MHz and 1 – 19 MHz. Average intensity in each channel is 0.5 photons/sample. (b) DFT spectra of the simulated waveform (blue trace in (a)) computed from 160 and 8,000 time samples. Note, the red plot is multiplied by 50. (c) Average DFT amplitudes at 7 MHz and 21 MHz computed from 400 simulated waveforms, representing comparison of signal and background. The corresponding waveform example is shown in (a) as the blue trace. Theoretical predictions from Eqs. (9) and (10) are shown as solid lines.

To illustrate adverse effects of cross-talk on image quality and measurement accuracy of functional signals, we created images from 1 to 10 combined waveforms, where each pixel intensity corresponded to *R_k_*(*f*) values (*f*= 1 – 19 MHz) in the DFT spectrum normalized to the expected intensity in units of photons/pulse. Only 160 time samples from each waveform with average amplitude of 0.5 photon/sample were used in computing pixel intensity. Here, waveform recording time per pixel was 1 μs, and signal intensity was equivalent to 1 photon/pulse, assuming 80 MHz laser repetition rate. Images shown in Fig. 3(a) are arranged in a 10 × 10 matrix. Rows contain images from 1 to 10 active channels each modulated at different frequency, and columns correspond to the specific modulation frequency One can see the progressive increase of noise in the images and the appearance of background as the number of active channels increases.

**Fig. 3.**
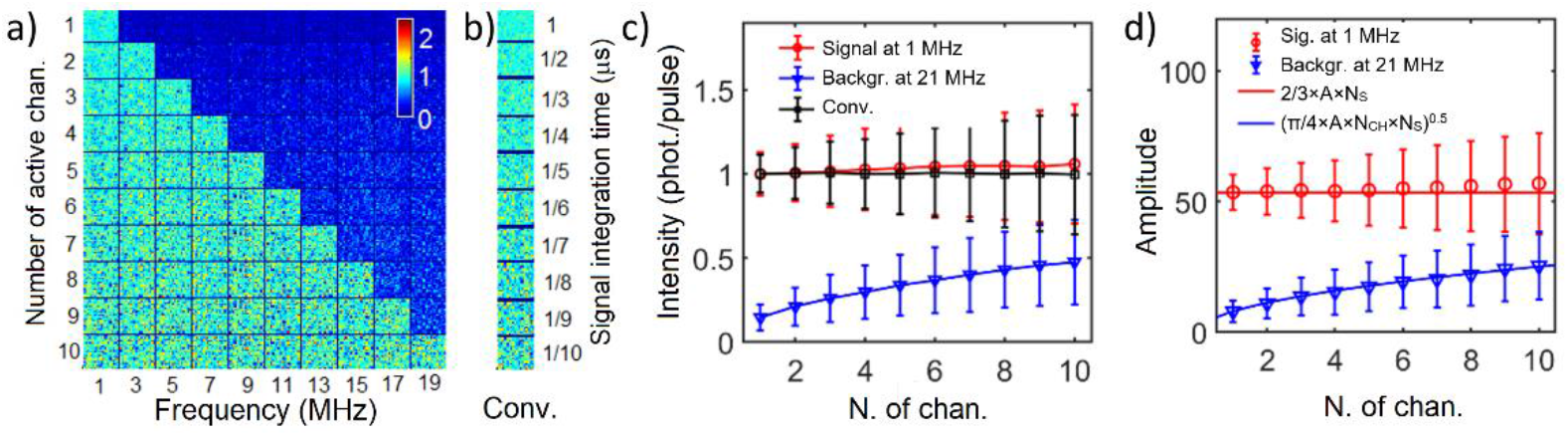
(a) Simulated frequency-multiplexed images from 1 to 10 active channels. False color intensity scale represents photons/pulse. (b) Conventional TPLSM images generated with different pixel dwell times. (c) Average and standard deviation of pixel intensity values computed from images in (a) and (b). (d) A comparison of theoretical estimates from Eqs. (9) and (10), and the data from (c) plotted as DFT amplitudes.

In our prior work [17], we discussed that frequency-multiplexed imaging performance in terms of SNR is similar to conventional TPLSM, given the same data acquisition rate and the same average signal intensity. Figure 3(b) shows simulated images from a conventional TPLSM that can be compared row by row to the images in Fig. 3(a). As the number of channels in 2P-FDM increases, the signal integration time in TPLSM is reduced proportionally. Pixel averages and corresponding standard deviations from images in Figs. 3(a,b) are presented in Fig. 3(c). There is a clear quantitative similarity in SNR values of TPLSM and 2P-FDM within the stated comparison criteria. Besides additional noise, each 2P-FDM image will contain ghost images from other channels due to shot noise-induced cross-talk, which will increase proportionally to the square root of sum of average signal intensity in every channel. In Fig. 3(d) we replotted the 2P-FDM data from Fig. 3(c) in units of DFT amplitude to demonstrate quantitative agreement with Eqs. (9) and (10).

Feasibility of using 2P-FDM for measurements of neural activity was explored with numerical analysis. We created simulations mimicking frequency-multiplexed recording of 10 calcium signals. In live experiments, signals from a neuron soma are typically acquired within 10-40 μs, depending on the speed of the scanner. The temporal resolution of calcium signals is defined by the frame acquisition time, which is much longer. Hence, we may neglect variations of calcium signal intensity within a single frame. First we synthesized 10 noiseless time-dependent reference signals which contained multiple spikes with similar kinetics, background level, and a range of amplitudes as seen in real calcium recordings. Each data point in these traces represented an average signal intensity expressed in units of photons/pulse. For every data point in each trace, a waveform containing 3,200 samples modulated at a single frequency was randomly generated, using the same set of frequencies as before. For each time point, the DFT of the sum of 10 corresponding waveforms was computed to obtain amplitude spectrum values at selected frequencies. Based on the same 10 reference traces, we generated datasets corresponding to calcium signals recorded with TPLSM. Similarly, for each data point defining the average signal intensity, we generated 320 samples and averaged them to obtain an estimate of a calcium signal amplitude in this measurement. Note, in 2P-FDM all signals are measured simultaneously within 20 μs, while in TPLSM these signals are measured sequentially within 2 μs each. Hence, both methods were compared within the stated criteria, i.e. same average signal intensity per channel and the same data acquisition rate. The results of this analysis are presented in Fig. 4. Figure 4(a) shows variations of a total signal computed as the sum of 10 reference signals, along with the sum of average number of photons of each corresponding waveform in 2P-FDM imaging and TPLSM in every frame. Signals are scaled in units of photons/pulse. Figures 4(b)–(d) show expected and computed signals from channels 1, 4 and 8, respectively. These plots represent 3 qualitative groups of calcium traces, starting from those with high intensity peaks and low background, and finishing with low intensity peaks comparable to the background level. One can see quantitative similarity of the results in 2P-FDM imaging and conventional TPLSM.

**Fig. 4.**
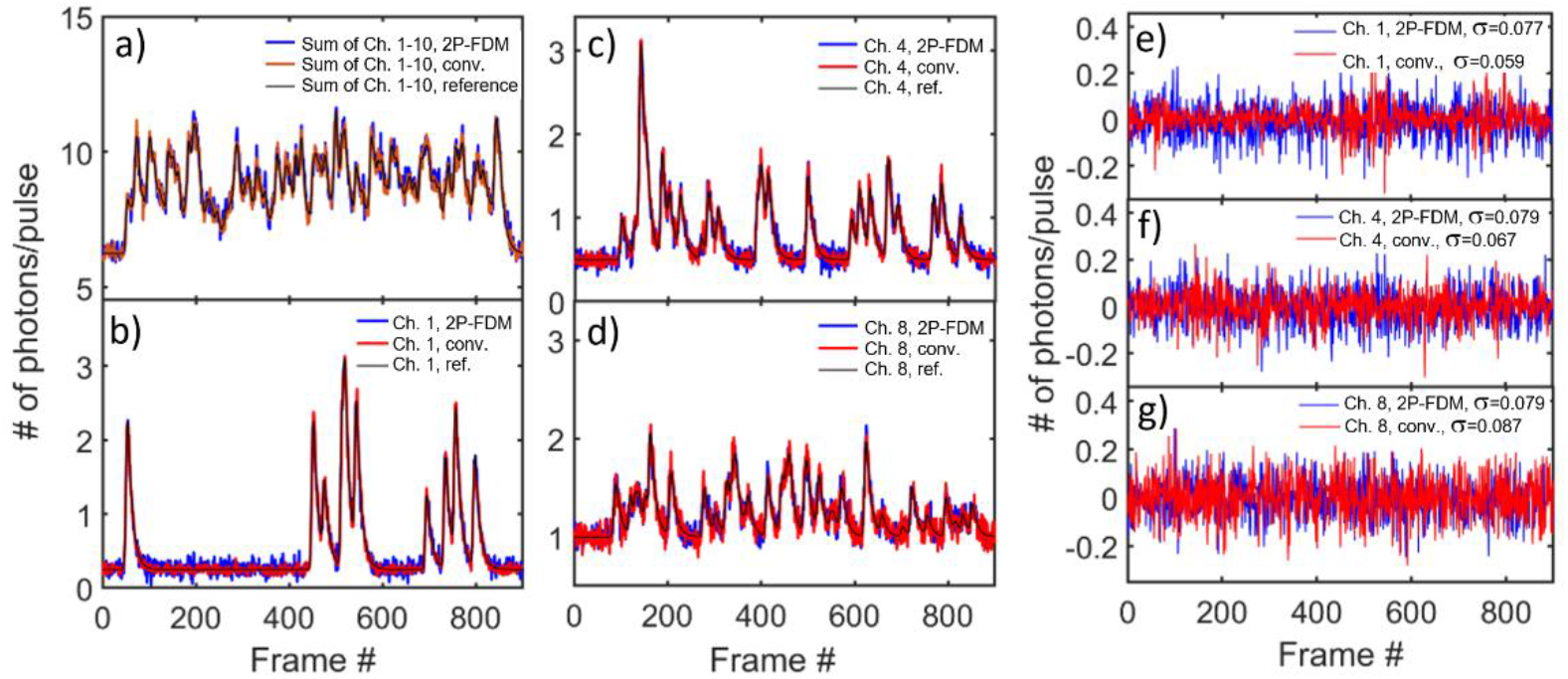
(a) Temporal profiles of simulated signals from all 10 channels. (b,c,d) Comparison of demultiplexed signals from channels 1, 4, 8 (*f* = 1,7,15 MHz), and conventional TPLSM signals. (e,f,g) Residuals in simulated 2P-FDM and TPLSM calcium signal traces computed from data in (b,c,d).

## 5. Frequency-multiplexed imaging *in vitro*

We acquired images from brain slices containing neurons labeled with the calcium indicator GCaMP6f by scanning a single neuron with 3 beams, modulated at 1, 5, 7 MHz (Fig. 3). The total excitation power at the objective was ~ 80 mW. For visualization purposes, the corresponding focal points in the image plane were laterally separated by ~ 10 μm along the horizontal axis. A high-speed digitizer with data streaming capabilities allowed acquisition of complete waveforms corresponding to scan lines across an image. Each line contained 1,274,000 samples, which were binned into 1,000 time samples/pixel during post-processing.

Figure 5(a) shows a sub-region of a conventional TPLSM image computed from the waveform/pixel average containing a triple image of the same neuron. The differences in image intensities of the neuron arise from small differences in power levels of the excitation beams and the nonlinear nature of the two-photon process. Besides the main three images, one can notice additional faint images of the same neuron (intensity < 10% relative to the main three) resulting from additional weak excitation beams. These extra beams are generated due to a non-linear response of the RF amplifier to periodic spikes in the amplitude of an AOD driving signal created by linear combination of three harmonics. These artifacts are present in the conventional two-photon image only, and not visible in the demultiplexed images due to low signal intensity. Waveforms demonstrating sections of a scan line containing the fluorescence signal of the neuron are shown in Figs. 5(b,c). The waveform in Fig. 5(c) contains ~30-50 distinct individual peaks within a 1 μs time window (*i.e*., 80 excitation pulses). Considering the possibility that higher intensity peaks could contain signals from multiple photons, we tentatively estimated a fluorescence signal intensity of ~1-2 photons/pulse in these data.

**Fig. 5.**
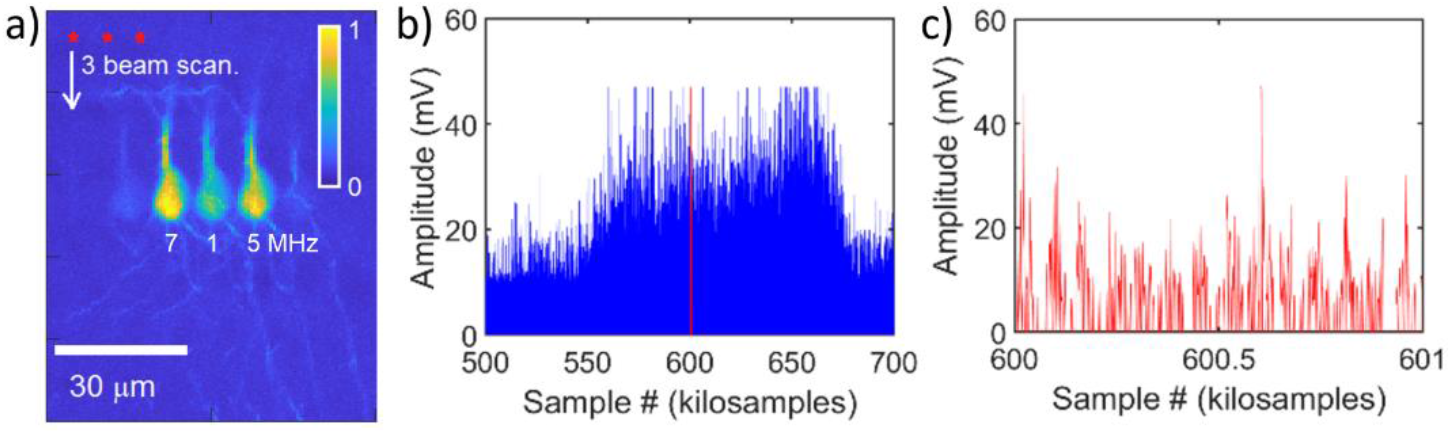
(a) Conventional two-photon image of a brain slice recorded while scanning the sample with 3 excitation beams, modulated at 7 MHz, 1 MHz, and 5 MHz. The image area shown contains 400 × 150 pixels along the vertical and horizontal axis, respectively. (b) A waveform section of a single scan line across the neuron on the left. The part shown in red represents 1,000 samples corresponding to 1 pixel in (a). (c) Waveform with 1,000 samples from (b) showing a fluorescence signal modulated at 7 MHz.

Demultiplexed imaging corresponding to different modulation frequencies are shown in Fig. 6. The first three images each exhibit a single neuron with high brightness along with two weak ghost images, which appear due to the wideband nature of shot noise in active channels modulated at different frequencies. The cross-talk increased with reduced pixel dwell time, as seen in the last three images. The image computed with 100 samples/pixel at 5 MHz shows almost no difference between neuron images from different frequency channels. Here, the pixel dwell time is only 0.2 μs, which is comparable to that of conventional TPLSM with fast resonant galvanometer scanning.

**Fig. 6.**
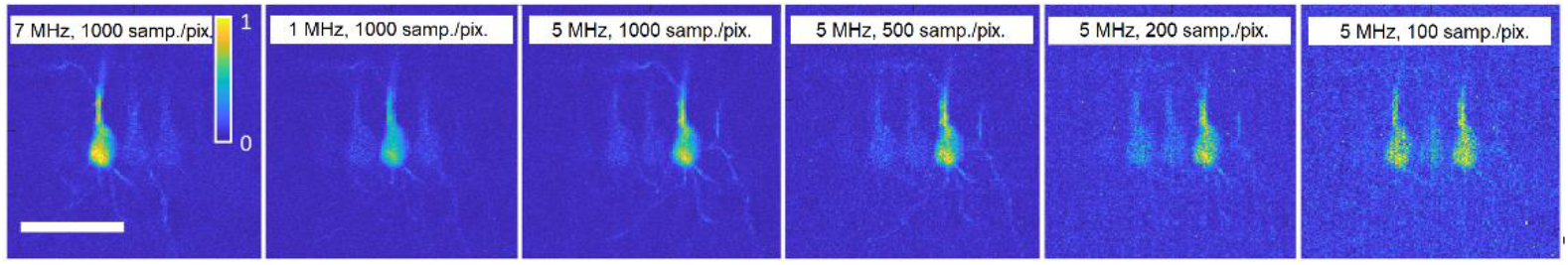
Demultiplexed images corresponding to different modulation frequencies and different number of samples/pixel as indicated. The scale bar is 30 μm.

Importantly, each pixel in these images contains inherent phase information, which we can extract by computing the 4-quadrant inverse tangent of the imaginary to real part ratio of the complex DFT value at a selected frequency. Figure 7 shows computed phase values from each pixel/waveform within identical ROIs, outlining cell bodies from Fig. 4(a). An example of cross-sections of the phase maps in Fig. 7(a) is presented in Fig.7(b). We observe relatively slow and gradual phase changes in all 3 images corresponding to different modulation frequencies, with additional variations superimposed on the general trend. A detailed analysis of such phase changes within 2P-FDM images in specific experimental conditions is the subject of a future study.

**Fig. 7.**
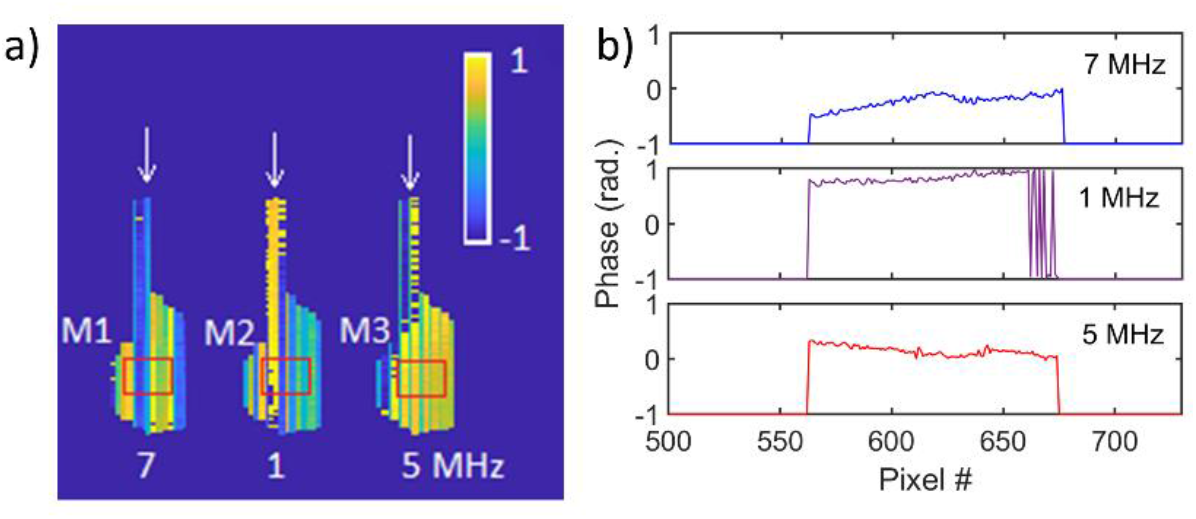
(a) Phase images computed from pixels within selected ROIs in Fig. 4(a) outlining cell bodies. (b) Cross-section plots from vertical lines in (a), indicated by arrows.

In frequency-multiplexed imaging, simultaneously generated signals from multiple sources are recorded with the same detector. In the current experimental configuration, the modulated beams were positioned very close, and thus they scanned essentially the same area. However, images from different frequency channels did not overlap, and only one channel was active at any given time. To simulate realistic conditions where multiple channels are active at the same time, we selected ROIs of 20×10 pixels from all 3 images of the neuron (red boxes M1, M2, M3 in Fig. 7(a)), and added the corresponding waveforms on a pixel-by-pixel basis. This selection was done to reduce the overall sample size and make it comparable to the number of samples that one may expect to acquire in conventional TPLSM from a single cell body. We created two datasets, [M1+M2] and [M1+M2+M3], consisting of 20×10× 1,000 samples and containing 2 and 3 amplitude-modulated signals, respectively. By progressively truncating the datasets from 1,000 down to 100 samples per waveform/pixel, we can analyze cross-talk effects between imaging channels as a function of pixel dwell time. Figure 8(a) shows examples of averaged DFT spectra computed from these datasets containing 1,000, 300, and 100 samples/pixel. In this analysis, the waveforms containing less than 1,000 samples/pixel were zero-padded to preserve the same set of frequencies in all DFT spectra. Active frequency channels are clearly visible in the top DFT plot. While one may expect a loss of spectral resolution with a decreased number of samples, the most important effect is the loss of contrast between the signal and the background. For example, DFT spectra computed from waveforms containing 100 actual samples show little difference between DFT spectra computed from both datasets, with and without 5 MHz modulation signal. In these conditions, frequency-multiplexed imaging will be severely impacted by the cross-talk. As follows from Eq. (11), cross-talk between channels in 2P-FDM can be reduced by binning the image into a smaller number of pixels while sacrificing spatial resolution.

**Fig. 8.**
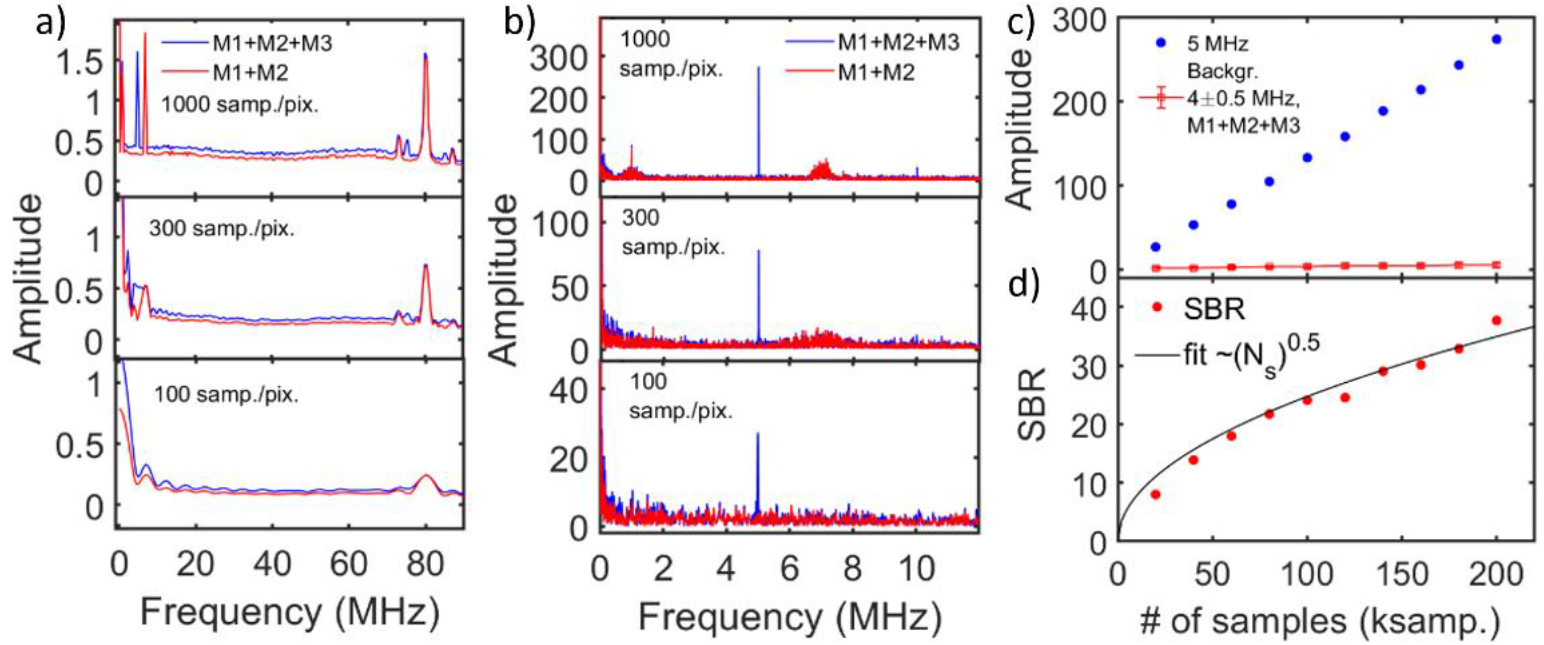
(a) Averaged DFT spectra of [M1+M2+M3] and [M1+M2] datasets computed with different number of samples/pixel. (b) DFT spectra of phase-aligned and concatenated waveforms from same datasets as in (a). (c) Signal at 5 MHz and average background computed from frequencies in the range of 3.5–4.5 MHz as a function of overall waveform length. (d) SBR plot computed from data in (c).

One of the most important applications of TPLSM is functional recording of fluorescence changes in calcium indicators linked to neural activity. The signal is usually recorded from a ROI occupied by neuron’s cell body which is more than 100× larger than the size of an individual pixel. To fully utilize advantages of the 2P-FDM method, it is essential to treat the signals acquired from a ROI as one continuous waveform which consists of individual parts corresponding to lines or pixels within the ROI. As we have explained above and demonstrated in Fig. 7, the phase of each pixel can be determined from the recorded trace itself. Other methods of tracking the phase not relying on phase calculation from noisy signals may be developed as well, *e.g*., by independently recording phases of both the excitation beams and the optical scanner. Whenever phase information is available, it is trivial to align the phases of each individual component and reconstruct an extended waveform. Specifically, we used phase information (Fig. 7(a)) corresponding to the specific frequency channel to adjust waveform phases to 0 by cyclic rotation of their samples (MATLAB’s *circshift* function). Due to waveform discretization, phase adjustments can be only performed with finite accuracy. The precision of phase adjustments may be improved, *e.g*., by interpolation and digital oversampling. Next, arrays were concatenated to form a single phase-aligned waveform. Figure 8(b) shows examples of DFT spectra of reconstructed waveforms corresponding to the 5 MHz channel obtained from the same datasets as in Fig. 8(a). Only the 5 MHz component is clearly visible in all spectra, while the remaining two frequencies are suppressed. Information from other frequency channels can be extracted in the same manner using the corresponding phase information (results not shown). We also observed that the order, in which pixels/waveforms are combined, affects the output DFT spectrum, especially for shorter waveforms. Because the observed phase of all waveforms varied relatively slow along a scan line, line by line concatenation maintained waveform correlation over multiple pixels for all active frequencies, and resulted in additional artifacts. Randomizing this steps, i.e. merging pixels/waveforms in a random sequence, will preserve the correlation for only a single selected frequency. The results shown in Fig. 8(b) were computed with such randomization. As we discussed earlier, in the presence of multiple active channels, the signal amplitude will be always measured against background spread across all frequencies. Figure 8(c) shows the DFT amplitude at 5 MHz and background values as a function of overall waveform length and the corresponding SBR plot, showing the expected square root of *N_s_* dependence.

## 6. TPLSM photon collection efficiency *in vivo*

Signal intensity is another critical parameter affecting cross-talk between frequency-modulated channels. To establish a connection of the numerical and theoretical results discussed earlier, we must quantify the number of photons detected in real experiments. One approach is measuring the system conversion gain *g* with the mean-variance or photon transfer curve method [20]. Although this approach is considered suitable for CCD and CMOS sensors, the method needs adaptation for amplified detectors, such as PMTs and avalanche photodiodes, due to the presence of additional noise sources increasing the signal variance [21]. To perform these measurements, another system was used, the Multiphoton Mesoscope [3], because of its highly optimized collection efficiency. To quantify *g*, we used a direct correlation of the average number of photons from a stable emission source measured by photon counting and the average pixel intensity from TPLSM images. In this experiment, waveforms showing individual photons and corresponding images of a fluorescein solution were acquired at different excitation power levels. Waveforms examples are presented in Fig. 9(a). Only spikes with amplitudes exceeding 40 mV were counted as detected photons. Figure 9(b) shows variations in the number of photons detected within a temporal window of 10 μs, and Fig. 9(c) shows the corresponding mean-variance plot. We observed a small deviation of ~ 15–20% from the expected trend, which may be caused by variations in the average photon flux as the beam moves across an imaging plane. Image non-uniformities in the range of ~ 10% are common in averaged TPLSM images recorded from fluorescein solution, and were reported previously [3]. Such variations do not significantly affect conversion gain measurements. In the central part of the scan line a resonant scanner moves with the highest speed, and with a 512 pixels and 12 kHz scanner the pixel dwell time is ~ 40 ns. The correspondence between average pixel intensity in the central part of the image and the average number of photons scaled proportionally to the pixel dwell time as shown in Fig. 9(d). Linear fit yielded a conversion gain of g ≈ 106 photons^−1^·pixel^−1^. We note, straightforward application of the mean-variance method without corrections yielded a noticeably smaller value of g ≈ 44 photons^−1^·pixel^−1^.

**Fig. 9.**
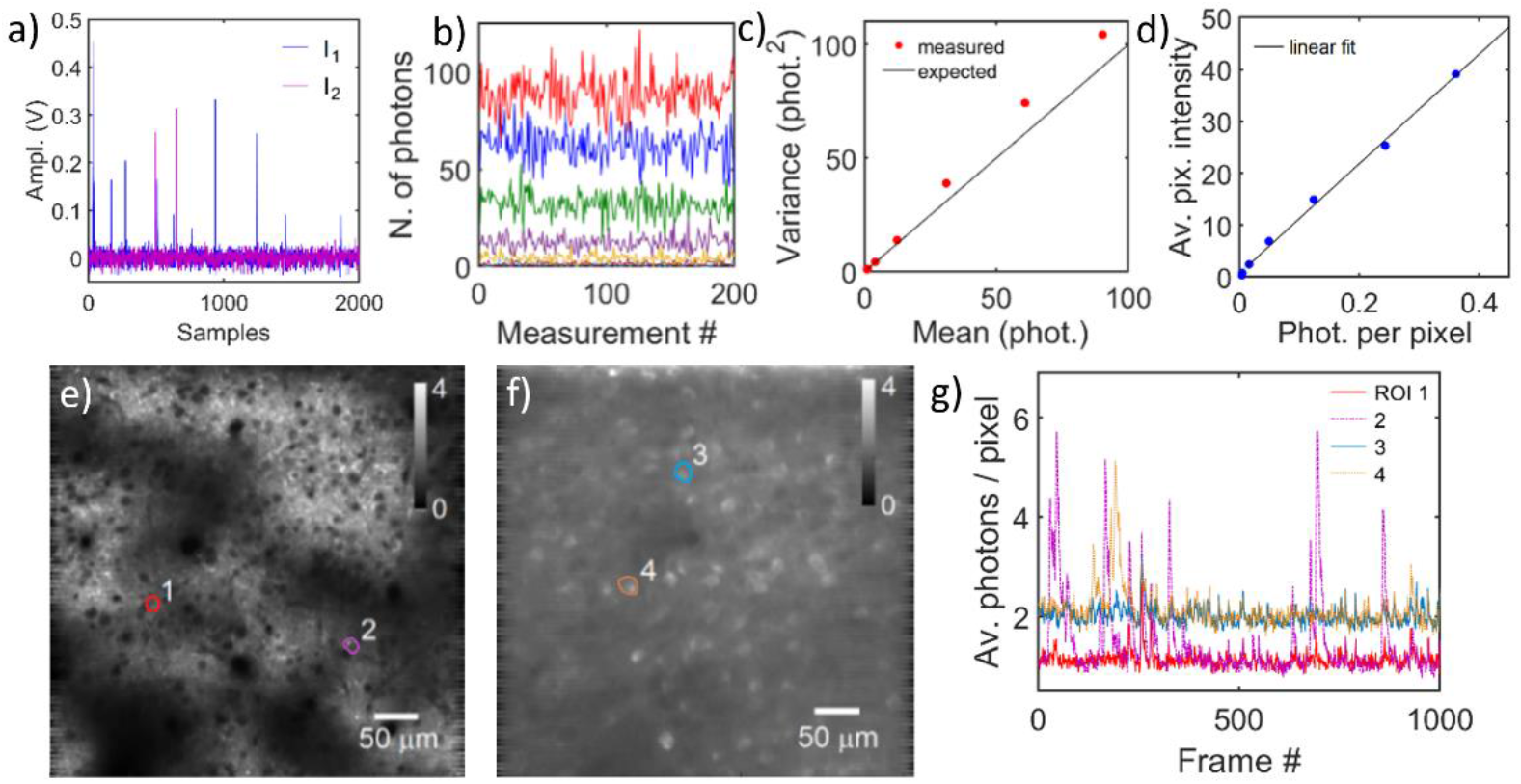
(a) Examples of PMT output showing individual photon events at different fluorescence intensities of a fluorescein solution. (b) Fluctuations in detected number of photons within a 10 μs temporal window, recorded at different excitation power levels. (c) Mean-variance plot of signals from (b). (d) Calibration plot showing average pixel intensity in TPLSM images as function of average number of detected photons. (e,f) Averaged *in vivo* images of mouse V1 layers 1 and 4 recorded at 75 μm and 375 μm depth. Intensity scale in units of photons/pixel. (g) Examples of calcium transients from selected ROIs in (e,f), recorded at ~ 18 Hz frame rate.

*In vivo* images of spontaneous activity within a mouse visual cortex recorded at depths of 75 μm and 375 μm, at the excitation power of ~ 70 mW and ~ 200 mW, respectively. The images averaged 1,000 times and scaled to show image intensity in units of photons/pixel, are presented in Figs. 9(e,f). Extracted calcium traces from selected cells scaled in units of average photons/pixel within the selected ROI are shown in Fig. 9(g). Each pixel integrates signals from 3-4 excitation pulses, and the observed fluorescence from selected cells vary in the range ~ 0.1 – 2 photons/pulse. Background signal fluctuations resulting from out-of-focus fluorescence appear at the level of ~ 0.1 photons/pulse (2 standard deviations).

## 7. Discussion

The major benefit of 2P-FDM is its ability to utilize a significantly higher number of imaging channels [15, 22] as compared to other multiplexed imaging methods, and to potentially achieve significant increases in imaging throughput as well as volumetric recording rate of functional brain activity. For example, a recent report utilized a similar concept to create continuous wave amplitude-modulated excitation to enable scanning with multiple beams, and demonstrated more than two orders increase in the imaging frame rate [23]. Unlike time-multiplexed approaches [12, 13, 16], 2P-FDM is not limited by the fluorescence lifetime of an optical indicator and one can utilize multiple laser sources without pulse synchronization between excitation channels. While our approach achieves high modulation frequencies which are needed to support high scanning speed, the maximum usable modulation frequency is ultimately limited by the laser pulse repetition rate and the pixel dwell time. To support separating individual frequency components, pixel dwell times must accommodate at least one, preferably several periods of each modulation frequency. For example, TPLSM with 512 pixels per scan line has a minimum pixel dwell time in the range of 40-120 ns, depending on the speed of the RG scanner, which corresponds to 3-10 excitation pulses per pixel at 80 MHz laser repetition rate. Thus, only the slowest RG scanners can support frequency-multiplexed imaging. However, the performance of 2P-FDM imaging can be improved by increasing the laser repetition rate, thereby increasing the range of usable modulation frequencies and the total amount of signal generated, as well as increasing pixel dwell time and binning the image into a smaller number of pixels.

Earlier, we discussed that 2P-FDM provides a SNR similar to conventional TPLSM at the same fluorescence signal intensity per channel and the same data acquisition rate [17]. This conclusion is further supported by the results of the numerical analysis, presented in Figs. 3(a,b). The important difference between these methods is that frequency-multiplexed imaging will always be affected by the cross-talk between different frequency channels due to the presence of shot noise. As follows from Eq. (11) (also see Fig. 3(c)), even with pixel dwell times of 1 μs and an average signal level of 1 photon/pulse, the SBR for 2 and respective 10 imaging channels will be approximately equal to 5 and 2. In this case, ~ 20% and 50% of the measured signal at a specific frequency will arise not from the channel of interest, but from all other imaging channels. However, with pixel dwell time of ~ 40 – 100 ns used conventional TPLSM the cross-talk levels will be significantly larger (see Fig. 6).

One of most significant applications of two-photon imaging technology is recording of functional information related to neural activity, reflected by dynamic calcium fluorescence signals from individual cells. In this analysis, only the signal averaged over multiple pixels corresponding to a neuron’s cell body is relevant. We proposed a new approach to analyze functional information recorded with the 2P-FDM method. Instead of signal averaging over multiple pixels, we treated all signals acquired from a cell body as one continuously acquired waveform. As we demonstrated in Fig. 9, the cross-talk contributions from other frequency-encoded channels become reduced proportionally to the square root of the acquisition time needed to measure a signal from every pixel within a given ROI. Note, the imaging speed is not reduced, and we only modified the method to process available experimental data. Since, the image of soma typically consists of more than 100 pixels, a more than 10-fold reduction in cross-talk levels between individual modulation channels can be expected with the newly proposed approach. One may use the theoretical description provided in this report to evaluate cross-talk levels in specific experimental conditions. For example, the signal acquisition time equals ~7 μs, given single cell body of ~15 μm in diameter and a dwell time of ~40 ns/μm when using the fastest 12 kHz RG scanner. Assuming average signal intensities of 0.5 photons/pulse in each imaging channel, the resulting SBR is approximately 9 and 6, for 2 and 4 imaging channels, respectively (see Eq. (11)). Therefore, we may expect to see noise at the level of ~ 0.05 – 0.08 photons/pulse when imaging 2 – 4 channels simultaneously. In the example shown above, the background noise of calcium signals appears at a similar level in the range of ~0.1 photons/pulse. Note, these measurements were performed with analog and not photon-counting detection, and therefore noise levels are slightly overestimated in this comparison. It is possible to conclude that at these fluorescence intensities and scan speeds 2P-FDM will allow reliable detection of calcium spikes from multiple simultaneously-imaged channels. Using slower RG scanners will even improve 2P-FDM performance. The results of numerical simulations presented in Fig. 5, which correspond to signal measurement time of 20 μs, also support feasibility of using the 2P-FDM method for functional recording of calcium activity.

A major limitation for 2P-FDM equally affecting every other TPLMS imaging method is the maximal physiologically acceptable excitation power for *in vivo* applications. Prior studies have placed this limit at ~250 mW with ~900 nm excitation [24]. Clearly, if multiple beam foci are distributed in the same imaging plane, the input power will increase proportionally to the number of excitation beams. Since the required power increases exponentially with depth, it appears advantageous to position the beams with axial offset and distribute laser power unevenly between scanning beams. Using a similar approach as in [25], where the excitation beam intensity is adjusted dynamically during imaging, depending on the localized signal intensity, it is possible to significantly reduce the overall exposure by restricting sample excitation only to the selected ROIs which include cell bodies and exclude regions that do not contain useful structures.

The following requirements must be met to allow the use of 2P-FDM for *in vivo* functional imaging. First, the scanning speed must be increased by using faster resonance galvanometer scanners. At the same time, methods to track phase of each amplitude-modulated excitation beam at high scan rates need to be developed. In this work we obtained phase information directly from the recorded emission signal. At high scan rates, reliable computation of each pixel’s phase will not be possible due to the presence of noise and the lack of spectral resolution between different frequency components. However, we expect that the required phase information can be determined from excitation waveforms, which can be recorded with high SNR, and possibly other variables such as phases of galvanometer scanners. Second, beam positioning is required to target different areas within an optically accessible imaging volume. Our current experimental setup can generate multiple frequency-encoded beams and simultaneously compensate for temporal and spatial dispersion of femtosecond pulses, yet it can only place excitation spots along a single line. Targeting specific areas is further hampered by the presence of harmonics that arise due to non-linear nature of the two-photon effect. One possible solution is to physically separate amplitude-modulated beams and adjust their position and focus with additional opto-mechanical components.

## 8. Conclusions

In this work, the utility of the 2P-FDM method for functional *in vivo* recordings of calcium signals was studied. Analytical expressions were derived, describing the relationship between average two-photon emission intensity in each channel and expected cross-talk arising due to shot noise in other active channels, and results were verified with numerical simulations. We proposed a novel approach that leverages phase information within individual pixels in the selected ROI to synthetically extend waveform measurement times and thus significantly reduce cross-talk between active channels. Practical application of this method was demonstrated *in vitro* by imaging GCaMP6f-labeled brain slices with multiple frequency-encoded beams. We also established a quantitative relationship between signal intensities detected during *in vivo* experiments and theoretical predictions to evaluate 2P-FDM performance. Our results suggest that calcium signal levels detected in real *in vivo* experiments are sufficient for 2P-FDM. Novel calcium indicators with increased fluorescence efficiency such as GCaMP7 [26] may further improve the performance of this imaging method. While we examined only one specific use case of 2P-FDM, other applications of this method are feasible. For example, spatially-multiplexed TPLSM with multiple excitation beams and multiple detectors [27] or random-access scanning [9] may incorporate frequency-encoded excitation and detection to improve data acquisition rates and the quality of recorded signals. We note that the 2P-FDM method is technically complex and still under development. Flexible positioning of excitation beams within the FOV, management of significantly higher data streams as compared to conventional TPLSM, real-time image reconstruction methods, as well as new image segmentation and post-processing algorithms are needed to fully utilize the advantages of 2P-FDM.

## Acknowledgments

We would like to thank James Brockill for proofreading the manuscript. We wish to thank the founder of the Allen Institute for Brain Science, Paul G. Allen, for his vision, encouragement and support.

